# Intra-Striatal Dopaminergic Inter-Subject Covariance in Social Drinkers and Nontreatment-Seeking Alcohol Use Disorder Participants

**DOI:** 10.1101/2024.02.06.579194

**Authors:** Evgeny J. Chumin, Mario Dzemidzic, Karmen K. Yoder

**Affiliations:** Psychological and Brain Sciences, Indiana University, Bloomington, IN, USA; Radiology and Imaging Sciences, Indiana University School of Medicine, Indianapolis, IN, USA; Stark Neurosciences Research Institute, Indiana University School of Medicine, Indianapolis, IN, USA; Neurology, Indiana University School of Medicine, Indianapolis, IN, USA

**Keywords:** PET, dopamine, striatum, network neuroscience, covariance networks

## Abstract

One of the neurobiological correlates of alcohol use disorder (AUD) is the disruption of striatal dopaminergic function. While regional differences in dopamine (DA) function have been well studied, inter-regional relationships (represented as inter-subject covariance) have not been investigated and may offer a novel avenue for understanding DA function.

Positron emission tomography (PET) data with [^11^C]raclopride in 22 social drinking controls and 17 AUD participants were used to generate group-level striatal covariance (partial Pearson correlation) networks, which were compared edgewise, also comparing global network metrics and community structure. An exploratory analysis examined the impact of tobacco cigarette use status. Striatal covariance was validated in an independent publicly available [^18^F]fallypride PET sample of healthy volunteers.

Striatal covariance of control participants from both datasets showed a clear bipartition of the network into two distinct communities, one in the anterior and another in the posterior striatum. This organization was disrupted in the AUD participant network, with significantly lower network metrics in AUD compared to the control network. Stratification by cigarette use suggests differential consequences on group covariance networks.

This work demonstrates that network neuroscience can quantify group differences in striatal DA and that its inter-regional interactions offer new insight into the consequences of AUD.

## Introduction

Disruption of striatal function and motivational circuitry is thought to underlie substance abuse and addiction, and dopamine (DA)-related alterations are believed to play a primary role (Schultz 2016). Despite being the most consumed substance of abuse, neurobiological mechanisms underpinning the transition to maladaptive alcohol use are not yet fully understood (Jordan and Andersen 2017; Nguyen-Louie et al. 2018; Volkow and Fowler 2000). Human neuroimaging studies of dopaminergic (DAergic) function employing positron emission tomography (PET) have reported altered DA function in various alcohol use disorder (AUD) populations (Martinez et al. 2005; Volkow et al. 1996; Volkow et al. 2017; Yoder et al. 2016). It has been proposed that alcohol use disorder is characterized by a reduction in DAergic function (although see Yoder et al. (2016)). Yet, we know little about whether regional interrelationships within the striatal DA system contribute to AUD.

The striatum is structurally interconnected with all major cortical structures, forming a topographically organized circuit loop that involves regulation of motor, sensory, and cognitive processing (e.g., motivation, reward, action planning) (Choi et al. 2017; Haber 2016). It is commonly subdivided into the caudate, putamen, and ventral striatum (comprised of the nucleus accumbens and ventral portions of anterior caudate and putamen) (Mawlawi et al. 2001). DAergic inputs onto the striatum originate from the substantia nigra pars compacta and ventral tegmental area in the brainstem. Previous work has shown that the striatum, although comprised of cytoarchitecturally overlapping areas of structural input, can be parcellated into spatially distinct regions of interest based on structural (Tziortzi et al. 2014) as well as functional connectivity (Tian et al. 2020). In the former study, Tziortzi et al. (2014) showed that homogeneity of DA D_2_ receptor availability (binding potential (BP)) was greater within the connectivity-based parcellation relative to commonly used anatomical divisions. Therefore, if regional connectivity is influencing the observed PET DA BP, regional DAergic function within the striatum is likely interrelated and could be investigated from a network perspective.

Unlike previous approaches that mostly focused on isolated brain areas, network neuroscience aims to study the brain as a system of interacting elements where each brain region is referred to as a node, and edges describe the connectivity between them. Edges can be quantified as absence/presence (e.g., binary structural connectivity), or by assessing their strength (i.e., weight), which could be streamline density for structural connectivity or a similarity measure such as correlation for functional connectivity (Rubinov and Sporns 2010). Additionally, given a homogenous sample/group of participants, it is also informative to compute similarity of inter-subject covariance (Veronese et al. 2019). Group differences in inter-subject covariance have been assessed for gray matter (Chang et al. 2018; Vijayakumar et al. 2021), metabolic activity ([^18^F]fluorodeoxyglucose PET) (Horwitz et al. 1984; Yakushev et al. 2017), and amyloid and tau-PET levels in Alzheimer’s disease (Franzmeier et al. 2019; Gonzalez-Escamilla et al. 2021). To our knowledge, there has only been one study on DAergic covariance, which investigated extrastriatal BP of [^11^C]FLB-457 PET in Parkinson’s disease (Mihaescu et al. 2021).

In this study, [^11^C]raclopride PET data were utilized to generate group level striatal covariance networks in social drinking controls (CON) and community recruited, currently drinking individuals with alcohol use disorder (AUD). Networks were then compared using graph theory metrics of density, degree, strength, clustering, and modularity (Rubinov and Sporns 2010). We hypothesized that covariance structure of the AUD group network would be altered as a function of chronic alcohol misuse. Additional exploratory comparisons were carried out to ascertain the potential impact of comorbid cigarette use on DA covariance in the striatum, as it has been shown that cigarette use alters DA function (Albrecht et al. 2013; Cosgrove et al. 2014; Zakiniaeiz et al. 2023). Finally, we used an open dataset of healthy controls with [^18^F]fallypride (FAL) DA PET imaging (Castrellon et al. 2019b) to replicate the observed striatal covariance structure.

## Methods

All participants provided written informed consent for all study procedures that were approved by the Indiana University Institutional Review Board in accordance with the Belmont Report. Participants were recruited via advertisement in local media, were 21-55 years old, and able to read, understand, and complete all study procedures in English. Exclusion criteria were history or presence of any psychiatric, neurological, or other medical disorder, current use of any psychotropic medication, positive urine pregnancy test, positive urine toxicology screen for illicit substances at interview, and contraindications for the magnetic resonance imaging (MRI). The Semi-Structured Assessment for the Genetics of Alcoholism was administered to confirm presence or absence of AUD (Bucholz et al. 1994). Participants also completed the following questionnaires: a medical history and demographics questionnaire, the 90-day Timeline Follow-Back for alcohol use (TLFB) (Sobell and Sobell 1992), Alcohol Dependence Scale (ADS), Fagerstrom Test for Nicotine Dependence (Pomerleau et al. 1994), Edinburgh Handedness Inventory (Oldfield 1971), AUD Identification test (AUDIT) (Saunders et al. 1993), and an internally-developed substance use questionnaire.

Demographic data for thirty-nine participants are summarized in Table 1. Education, biological sex, and smoking variables differed between AUD and CON groups, and as expected AUD scored significantly higher on the AUDIT. Six CON participants tested positive for illicit substances on scan day (5 marijuana, 1 benzodiazepine) as well as one AUD participant (marijuana) and were included in the study due to rarity of endorsed use.

**Table 1.** Demographic Data. Data are shown as mean ± standard deviation or number of participants. ^*^Denotes significant group differences at *p* < 0.05, using a two-sample *t*-test or *χ*2 test as appropriate.

|  | Group (n) | Age (years) | *Education (years) | *Sex (M/F) | Handedness (R / L / RL) | Race (W / AA / Other) | *Smoking | Fagerstrom | *AUDIT |
| --- | --- | --- | --- | --- | --- | --- | --- | --- | --- |
| Social drinking controls (CON) | 22 | 39.9 ± 10.8 | 15.8 ± 2.0 | 10/12 | 17/4/1 | 11/9/2 | 3 | 3 ± 0 | 6.7 ± 3.6 |
| Nontreatment-seeking alcohol use disorder (AUD) | 17 | 43.0 ± 10.1 | 13.5 ± 2.2 | 14/3 | 16/1/0 | 10/7/0 | 10 | 5 ± 1.7 | 15.2 ± 5.2 |

### Imaging

Structural magnetic resonance imaging was performed on a Siemens 3T SKYRA (10 controls and 8 AUD) and a Siemens 3T PRISMA (12 controls and 9 AUD), with a high resolution T1-weighted anatomical image acquired with a whole brain magnetization prepared rapid gradient echo (MPRAGE) sequence (1.05×1.05×1.2 mm^3^ voxels; 176 sagittal slices). PET data were acquired on a Siemens Biograph mCT scanner (Siemens, Erlangen, Germany). PET scans were initiated with a single bolus injection of 14.35 ± 0.92 mCi of the DA D_2_/D_3_ antagonist [^11^C]raclopride (RAC). A CT scan was acquired for attenuation correction. Dynamic acquisitions were 50 minutes in length, where the first 10 frames were 30 sec. long, while the remaining 45 frames were 60 sec. long. Data were reconstructed with Siemens software using the Filtered Backprojection algorithm (5 mm Hanning filter) with corrections for random coincidences, attenuation, and scattering.

### Structural Image Processing

Structural T1-weighted MRI data for each participant were first denoised (Coupé et al. 2008), bias field corrected and tissue-type segmented (*fsl_anat*, FSL v6.0 (Jenkinson et al. 2012)), and then skull stripped with ANTs (Tustison et al. 2014). A combination of linear (*flirt* 6 then 12 degrees of freedom) and nonlinear (*fnirt*) transformations in FSL was used to register T1 volumes to a standard space (Montreal Neurological Institute template), with the inverse transformations applied to align the Melbourne subcortical atlas Scale II parcellation (Tian et al. 2020) to each participant’s T1 image (Jenkinson et al. 2012).

### PET preprocessing

RAC PET data preprocessing was done in the Statistical Parametric Mapping software version 12 (SPM12; https://www.fil.ion.ucl.ac.uk/spm/software/spm12/). For each participant, the first 14 frames (∼10 min) were averaged to create an early mean PET image that was used as a target for realignment of individual frames to correct for head motion. The mean image was then coregistered to the structural MRI with the transformation applied to dynamic data. Subcortical regions of interest from the Tian et al. (2020) parcellation (left and right anterior and posterior caudate and putamen, and nucleus accumbens shell and core) were used as nodes in the inter-subject covariance network generation. An additional cerebellar gray matter (with the vermis excluded) reference region (with little or no D_2_/D_3_ binding) was defined in the native T1 space of each participant. Regional time activity curves for the striatal subregions and cerebellum were generated in Matlab (R2022b). DA tone was defined as RAC D_2_/D_3_ receptor availability (binding potential (BP_ND_); relative to nondisplaceable binding) (Innis et al. 2007). Regional BP_ND_ was estimated with the Multilinear Reference Tissue Model (MRTM; Ichise et al. (2003)).

### Network Construction

Inter-subject covariance networks for each group consisted of twelve striatal nodes described above with edges quantified as partial Pearson correlation coefficients, adjusting for age, biological sex, and smoking status of participants within group. This approach generates a signed fully connected adjacency matrix. For exploratory analysis of cigarette use, networks were created with partial Pearson correlation (age and sex adjusted) from three groups: nonsmoking CON (n=19), nonsmoking AUD (n=7), and smoking AUD (n=10). Additionally, sparse networks were computed by identifying edges of significant covariance via permutation testing (10,000 permutations) where a null distribution of networks was generated by randomly and independently scrambling the regional BP_ND_ values of each participant and recomputing the covariance matrix. This procedure destroys any existing regional inter-subject relationships. A permutation-based *p* < 0.05 value was then used to identify significant edges. Stricter significance thresholds resulted in fragmented networks in the AUD group, in which one or multiple regions became disconnected from the rest. Network fragmentation prevents direct group comparisons and limits interpretability, therefore, only the *p* < 0.05 thresholded networks are reported. To assess whether the covariance structure was specific to BP_ND_, the tissue-to-plasma rate constant *k*_2_ and tracer delivery *R*1 parameters from MRTM as well as region size (in voxels) were also used to generate group covariance networks.

### Fallypride Validation Dataset

Data from 25 healthy young adults (12 male and 13 female; age 18-24) as described in Castrellon et al. (2019b) were downloaded from OpenNeuro (Castrellon et al. 2019a). The provided standard space parametric images of simplified reference tissue model derived BP_ND_ were used to extract regional averages for striatal nodes described above. Networks were constructed with partial Pearson correlation, with age and education as covariates.

### Global and Nodal Metrics

The following measures were computed from group covariance networks as described in the Brain Connectivity Toolbox (Rubinov and Sporns 2010): density - number of connections [thresholded networks only], node degree – number of connections for each region [thresholded networks only], positive and negative strength – sum of edge weights (i.e., positive or negative correlation coefficients) for each region (summing over its connections) and for the whole network (summing all edges) [unthresholded and thresholded networks], and clustering coefficient – a measure of the tendency of nodes in a network to cluster (using the weighted and signed variant (Costantini and Perugini 2014)) [unthresholded and thresholded networks].

Community structure was estimated from each network with the multiresolution consensus clustering (MRCC) method (Jeub et al. 2018), with 10,000 initial partitions sweeping the resolution parameter space. A co-assignment matrix (probability that any region pair is assigned to the community) was then computed, followed by application of a null model for identification of a single consensus partition (here the alpha threshold for the null model was set at 0.05). For within community (a subset of regions grouped together by the MRCC algorithm) averaged BP_ND_ comparisons, the CON group consensus partition was a reference partition and applied to all groups.

### Statistical Analysis

Statistical comparisons were performed in Matlab 2022b. Demographic, regional BP_ND_, and community-averaged BP_ND_ comparisons were assessed via independent samples *t*-tests or *χ*^*2*^ tests as appropriate. Distributions of nodal network values were compared via a 2-sample Kolmogorov-Smirnov test. The three-group exploratory analysis applied a one-way ANOVA with Tukey-Kramer *post hoc* tests.

### Code Availability

Image processing and covariance network analysis scripts have been made publicly available: structural T1-weighted image preprocessing (https://github.com/IUSCA/IUSM-ConnPipe), RAC PET processing (https://github.com/echumin/PET_processing_Code), and Covariance network analysis (https://github.com/echumin/CovNet).

## Results

Among all striatal regions/nodes, only the left anterior caudate BP_ND_ was significantly lower in AUD compared to CON (Table 2; independent-samples *t*-test *p* = 0.009, uncorrected). However, this difference was not significant after accounting for multiple comparisons (false discovery rate (FDR) *q* < 0.05).

**Table 2.**
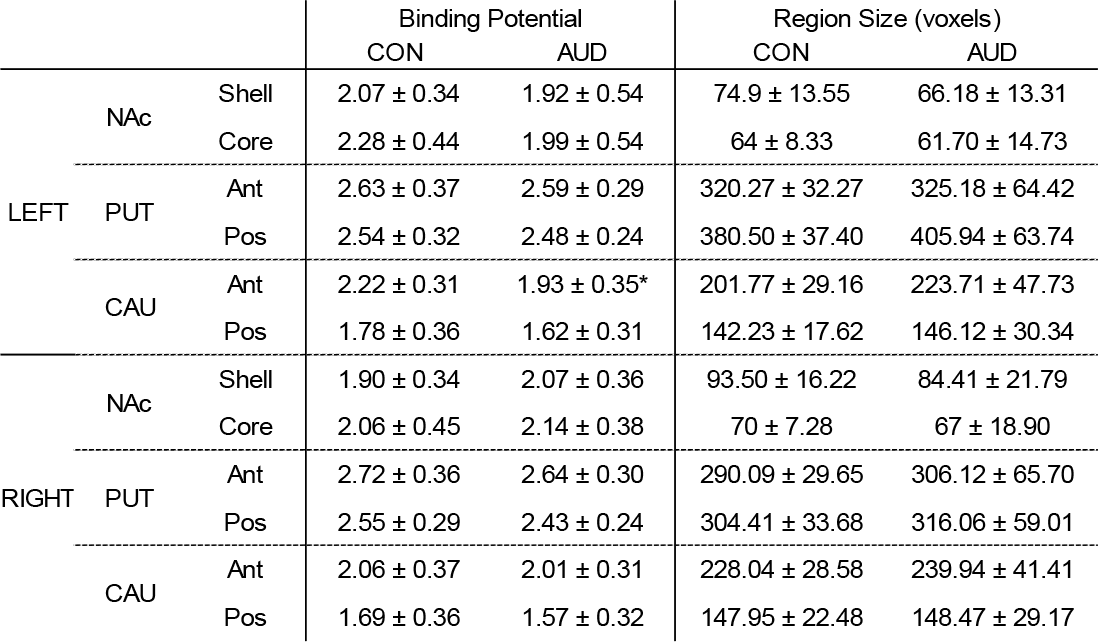
Regional Size and Binding Potentials. Mean ± standard deviation of binding potential and region size values (in voxels from 2 mm isotropic resolution volumes) for the twelve striatal regions of interest from the Scale II Melbourne subcortical atlas (Tian et. al., 2020). *Denotes group differences at *p* < 0.05, uncorrected independent samples *t*-test. Ant – anterior, Pos – posterior, NAc – nucleus accumbens, PUT – putamen, CAU – caudate, CON – controls, AUD – alcohol use disorder.

Within participant Z-scored RAC BP_ND_ values for each node are shown in Figure 1A and 1D for CON and AUD, respectively. The covariance network of the CON group showed a clear pattern of similarity among bilateral nucleus accumbens (shell and core) and anterior putamen that was anticorrelated with a separate set of regions comprised of bilateral caudate (anterior and posterior) and posterior putamen (Figure 1B). This pattern was less evident in the AUD group network (Figure 1E). Edge weight distributions between the two group networks showed no differences (KS=0.18, *p*=0.2). Permutation significance-based thresholding of correlations (edge weights) revealed a greater number of edges in CON relative to AUD (Figure 1C and 1F; network densities 0.73 and 0.42 for CON and AUD groups, respectively). To assess whether the observed covariance structure was specific to BP_ND_, we also computed networks of MRTM flow constants *k*_2_ and *R*1 as well as region size. These MRTM flow constant networks did not show the organization seen with BP_ND_ (Supplementary Figure 1; no group differences were found for *k*_2_ or *R*1 for any region), while volumetric covariance showed a pattern distinct from BP_ND_ covariance, displaying positive covariance among bilateral nucleus accumbens and putamen and a separate group comprised of bilateral caudate regions (Supplementary Figure 2).

**Figure 1.**
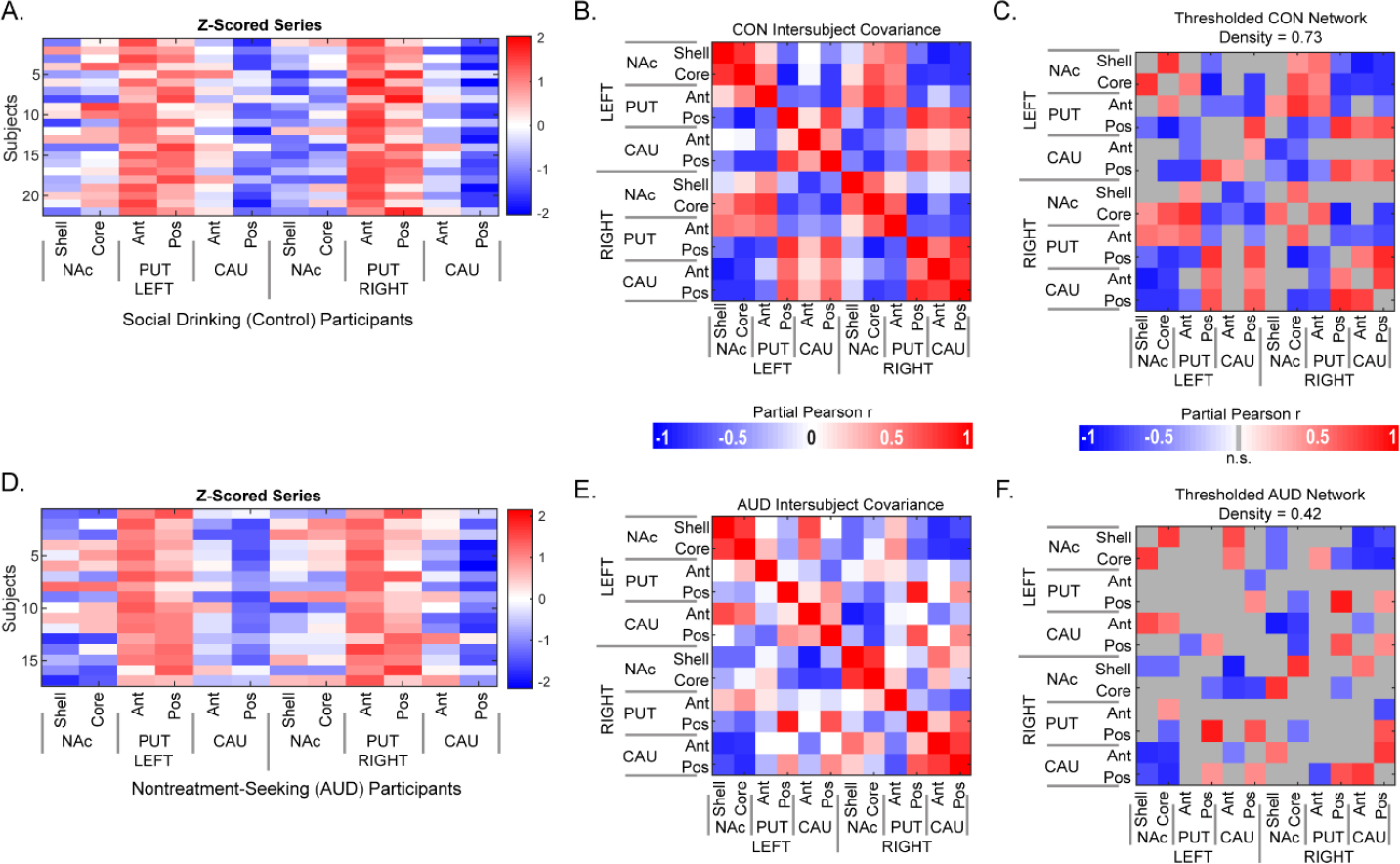
Inter-subject group covariance of striatal [^11^C]raclopride binding potential. Subject-series of RAC BPND for striatal regions of interest are shown for **(A)** control and **(D)** AUD participants. Covariance matrices **(B, E)** were computed as pairwise partial Pearson correlation and adjusted for age, biological sex, and cigarette smoking status. Permutation testing identified edges of significant covariance **(C, F)** within each group (*p* < 0.05). Density values show fraction of nonzero edges in the network. Ant – anterior, Pos – posterior, NAc – nucleus accumbens, PUT – putamen, CAU – caudate, CON – controls, AUD – alcohol use disorder.

Elementwise (edge-level) group comparisons of the two networks identified three edges at uncorrected permutation *p*<0.05 significance: (1) left nucleus accumbens shell and anterior caudate, (2) left nucleus accumbens core and right nucleus accumbens shell, and (3) left and right nucleus accumbens core covariances (Figure 2A). In addition, total positive and negative strengths were lower in the AUD for both unthresholded and thresholded networks (Figure 2B-C). AUD also had lower nodal measures of degree, clustering coefficient, and positive and negative nodal strengths (Figure 2D-G).

**Figure 2.**
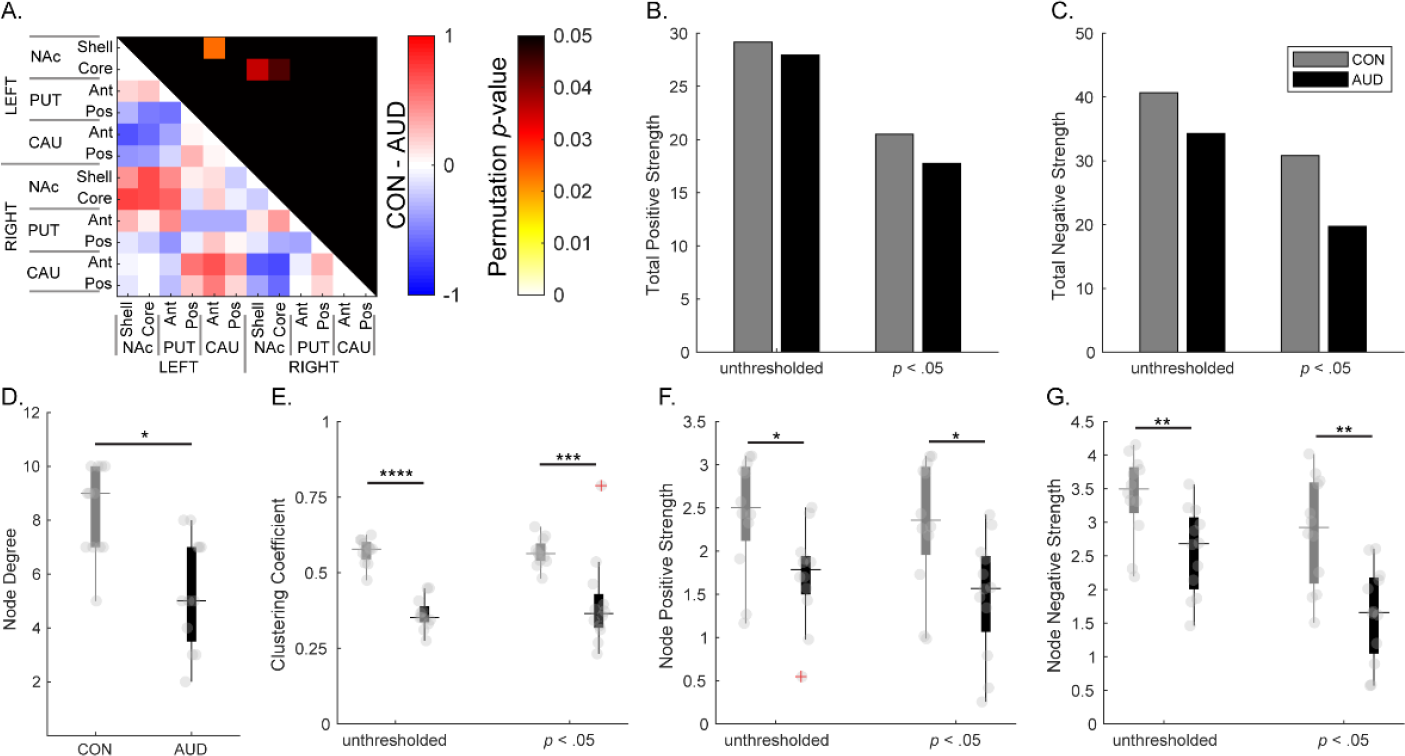
Group comparisons of CON and AUD covariance networks. **(A)** Covariance differences of weights for CON minus AUD matrix (lower triangle) and permutation test *p-*values (uncorrected) of edge-level significant group differences (upper triangle). **(B)** Positive and **(C)** negative total strength values of covariance networks (unthresholded and *p* < 0.05 correlation significance thresholded). **(D-G)** Distributions of nodal measures of covariance networks: **(D)** node degree, **(E)** clustering coefficient, and **(F)** positive and **(G)** negative nodal strengths. Differences in distributions from the two networks were assessed with a two-tailed Kolmogorov-Smirnov test. ^*^ *p* < 0.05, ^**^ *p* < 0.005, ^***^ *p* < 0.001, ^****^ *p* < 0.00001. Ant – anterior, Pos – posterior, NAc – nucleus accumbens, PUT – putamen, CAU – caudate, CON – controls, AUD – alcohol use disorder.

Community detection identified a two-community partition for each group. For the CON covariance network, the partition grouped bilateral nucleus accumbens and anterior putamen into one community and bilateral caudate and posterior putamen into another (Figure 3A). These communities were clearly visible in the correlation matrix, even without employing community detection. The covariance structure in the AUD included one large community and a second smaller community that was comprised of bilateral posterior putamen and left posterior caudate (Figure 3B, lower triangle). From the initial set of 10,000 partitions the probability of any region pair being assigned to the same community can be expressed in a co-assignment matrix (Figure 3C). From the AUD co-assignment matrix, it is evident there is no clear community partition and that the homotopic similarity observed in CON is absent. The thresholded matrices for both groups (Figure 3D) showed no cross-hemisphere covariance of nucleus accumbens and anterior putamen in AUD that was observed in the CON network. Additionally, using the CON partition as a reference, we quantified differences in communities between groups by computing within and between module strengths as well as mean BP_ND_ in each community. For unthresholded networks, within community positive strengths (Figure 3E) and between community negative strengths (Figure 3F) were lower in AUD. When examining the average within module BP_ND_ for the two communities (Figure 3G); there were no significant group differences for either module.

**Figure 3.**
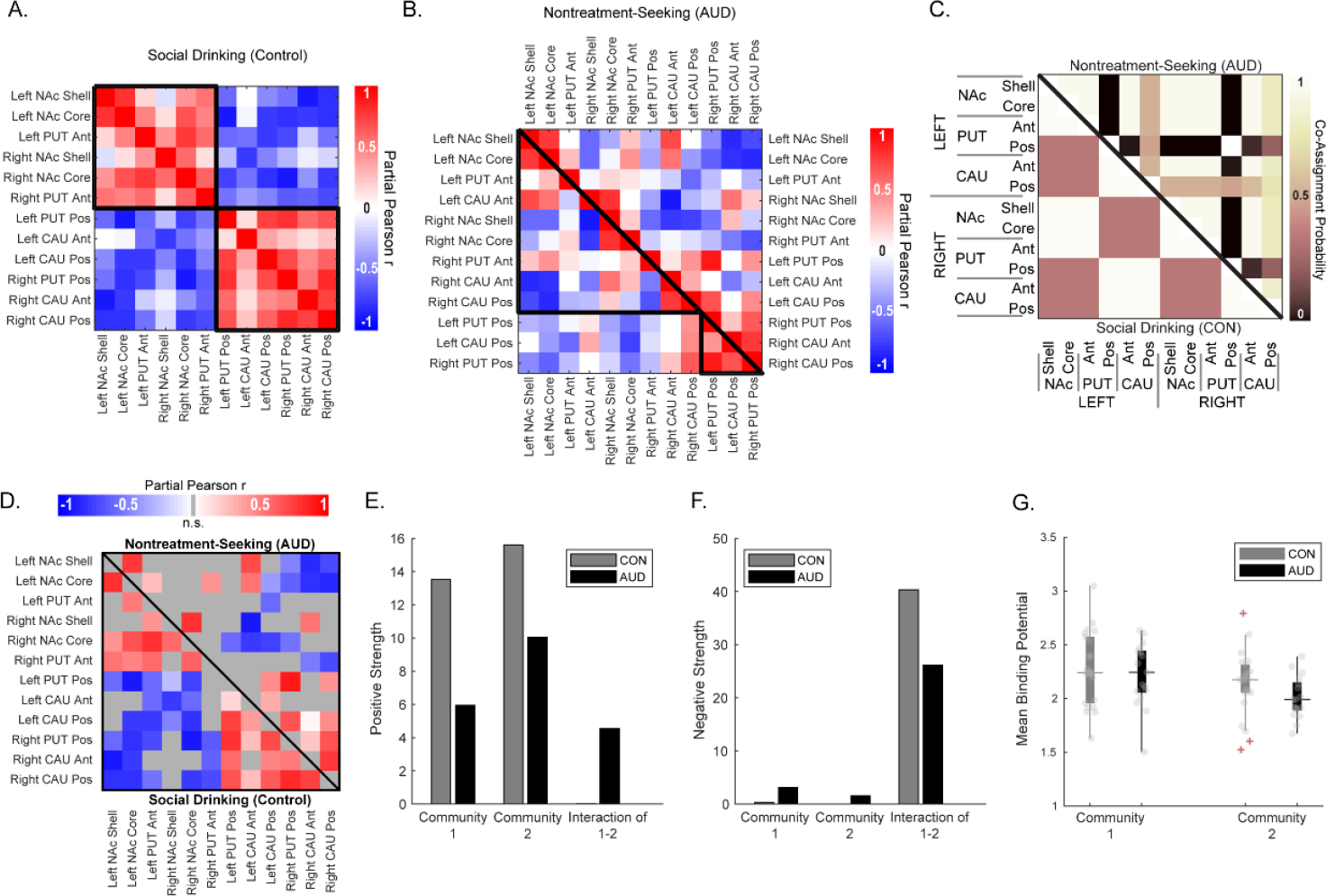
Community structure of inter-striatal covariance. **(A)** Social drinking controls (CON) network community detection revealed a bipartition of the network into two communities. **(B)** Nontreatment-seeking alcohol use disorder (AUD) covariance network was also a bipartition (lower triangle), however, these communities were distinct from CON, as seen in upper triangle where regions are arranged by community affiliations form the CON network. **(C)** Co-assignment matrices for CON (lower triangle) and AUD (upper triangle) show the probability of any two nodes being assigned to the same community from the initial ensemble of 10,000 partitions, illustrating the disrupted community structure in AUD relative to CON. **(D)** CON community structure ordered networks for CON (lower triangle) and AUD (upper triangle). **(E-F)** Total strengths within each community and the between community blocks of the matrices: **(E)** positive and **(F)** negative strengths. **(G)** Community average binding potentials were computed per participant and displayed by group. Ant – anterior, Pos – posterior, NAc – nucleus accumbens, PUT – putamen, CAU – caudate, CON – controls, AUD – alcohol use disorder.

### Exploratory Cross-Sectional Analysis of Cigarette Use

We stratified the sample on cigarette use and computed covariance networks for nonsmoking controls (nsCON; n=19), nonsmoking AUD (nsAUD; n=7), and smoking AUD (smAUD; n=10) subgroups. The three smoking controls were not sufficient to perform analysis in this group. Overall, we observed a similar pattern, where the AUD groups had altered covariance of homotopic regions, notably the nucleus accumbens, as well as reduced thresholded network densities (Figure 4A-C). Community structure analysis revealed the highest variance in community assignment was in smAUD (Figure 4D-F, left triangles). The loss of cross-hemisphere covariance is also observed in the community ordered matrices for each group (Figure 4D-F, right triangles), such that homotopic regions (e.g., the nucleus accumbens), no longer group into the same community within the AUD subgroups. Using nsCON partitions as reference, average BP_ND_ values were compared for each module across subgroups. Module 2 (comprised of bilateral caudate and posterior putamen) had a significantly lower BP_ND_ in smAUD relative to nsCON group (one-way ANOVA *F*(2,33) = 4.75, *p* = 0.015, with post hoc Tukey-Kramer tests, *p* = 0.011).

**Figure 4.**
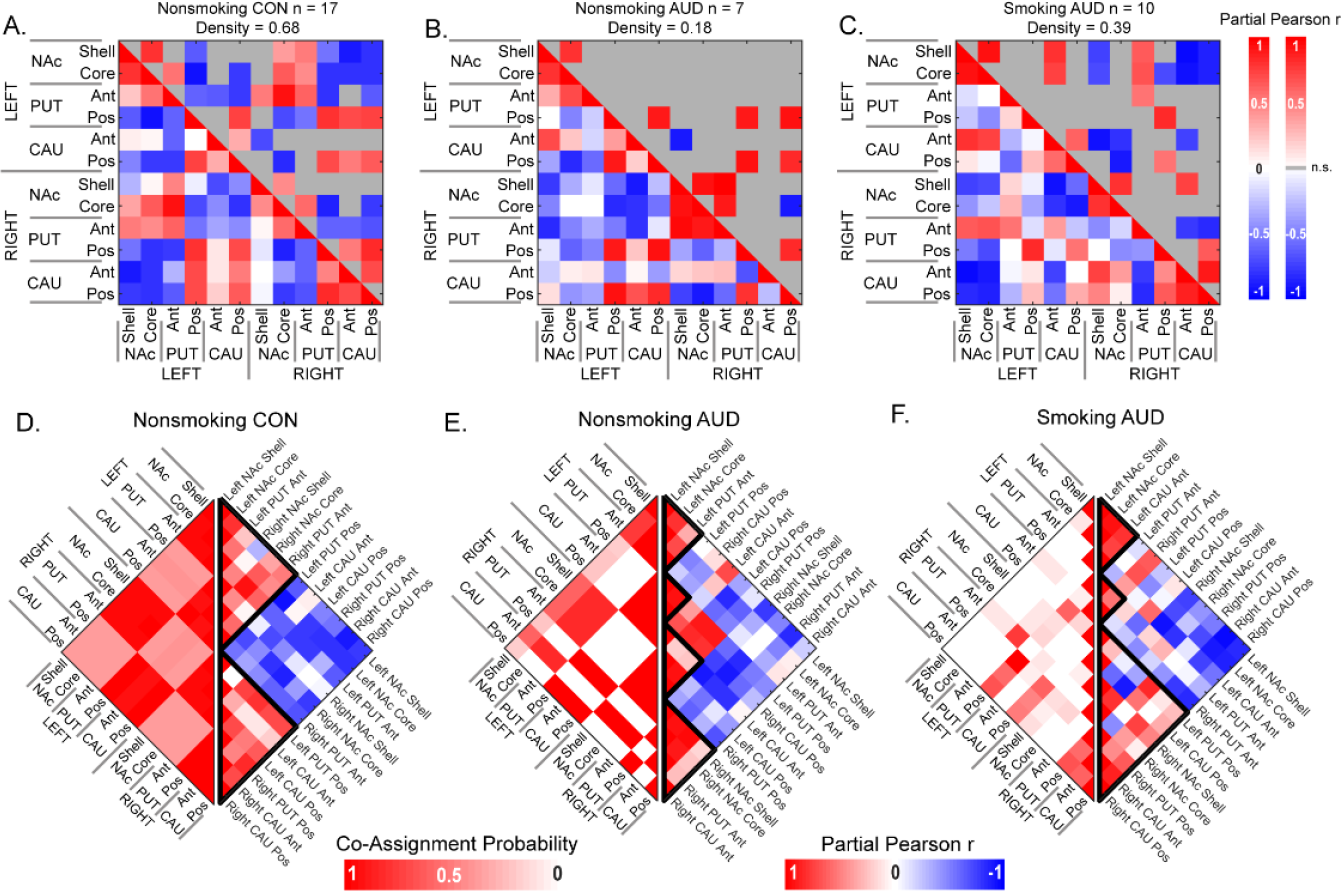
Covariance and community structure of nonsmoking control (CON; A, D), nonsmoking alcohol use disorder (AUD; B, E), and smoking AUD (C, F) subgroups. **(A-C)** Lower triangle matrices show the covariance networks (partial Pearson correlation, adjusted for age and sex) for **(A)** nonsmoking CON, **(B)** nonsmoking AUD, and **(C)** smoking AUD. Upper triangle matrices show significant edges (*p* < 0.05, permutation testing) for each covariance network with network densities (number of nonzero edges) reported above each matrix. **(D-F)** Multiresolution consensus clustering (MRCC) co-assignment matrices (left triangle), with their respective community ordered covariance matrices in the right triangles (same data as A-C (lower triangles) but reordered based on identified community structure for each group network). Ant – anterior, Pos – posterior, NAc – nucleus accumbens, PUT – putamen, CAU – caudate.

### Validation in the Independent [^18^F]Fallypride Dataset

We leveraged a publicly available FAL PET dataset (Castrellon et al. 2019a; b) to assess whether the striatal DAergic covariance observed in our RAC PET dataset was capturing DA receptor distribution specific variance. Comparison of FAL covariance and RAC covariance showed strong similarity (Figure 5), with a significant correlation of edge weights (Pearson’s *r* = 0.66, *p* < 5×10^−9^). Interestingly region pairs whose weights showed opposite relationship were predominantly interhemispheric (Figure 5C).

**Figure 5.**
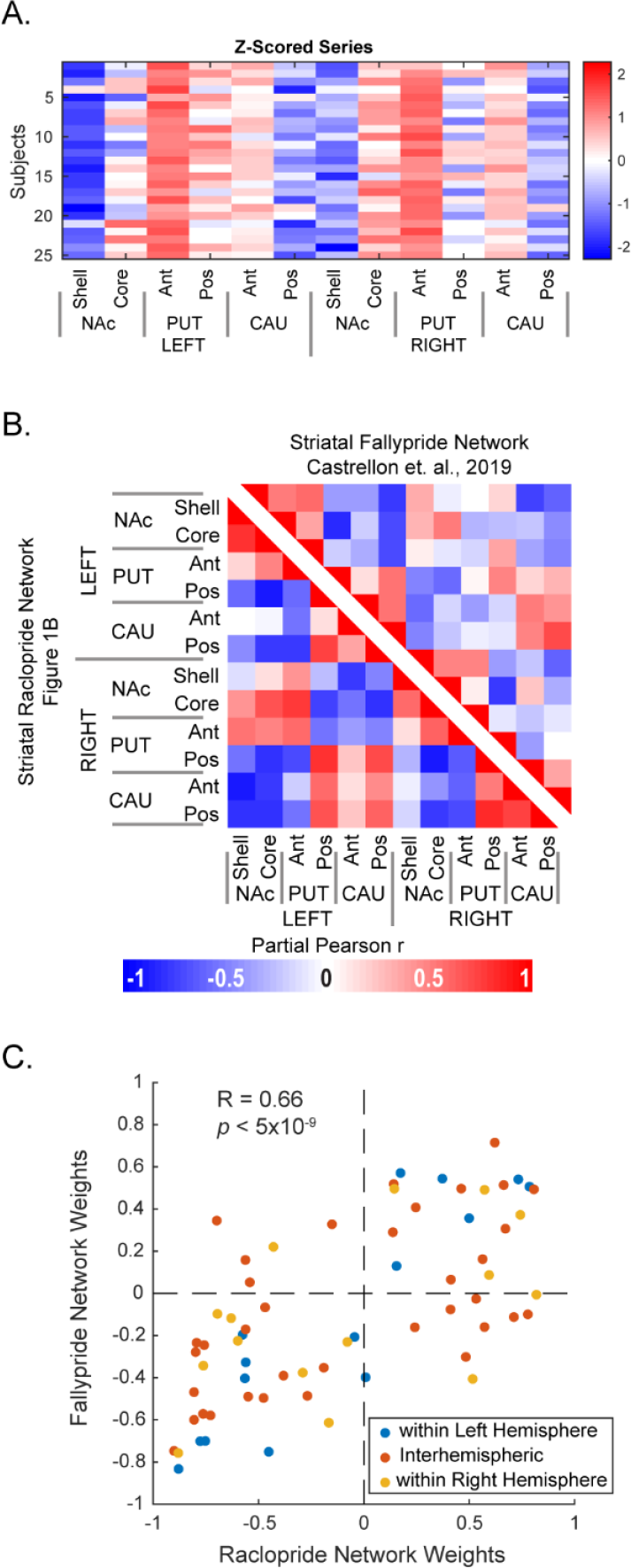
Comparison of inter-subject group covariance networks from [^11^C]raclopride and [^18^F]fallypride quantified striatal dopamine. **(A)** Subje series of fallypride Z-scored BPND for stri region of interest. **(B)** Side by side comparison of raclopride (lower triangle) fallypride (upper triangle) derived networ (partial Pearson correlation, adjusted for and sex). **(C)** Edgewise cross-correlation network weights between the raclopride fallypride derived networks presented in Color coding distinguishes edges within between hemispheres. Ant – anterior, Po posterior, NAc – nucleus accumbens, P putamen, CAU – caudate.

## Discussion

Here we show that network neuroscience methodology can successfully quantify the interrelationships of DAergic striatal function to study neural correlates of AUD. Structured covariance of RAC BP_ND_ observed among striatal regions of the social drinking CON group was disrupted in the active AUD group. This pattern of covariance was not present with networks generated from tracer kinetic model flow parameters k2 and R1 or from region size for either CON or AUD group (Supplemental Figures). This covariance structure was also replicated in an independent dataset from a publicly available FAL study of healthy volunteers from (Castrellon et al. 2019b). In sum, these results suggest that the striatal covariance structure we observed is not limited to our sample or the raclopride PET tracer, but rather captures biologically meaningful information on DAergic function in the striatum.

We used the Melbourne subcortical atlas to parcellate the striatum into twelve nodes, which can be anatomically aggregated into the caudate, putamen, and ventral striatum (which includes the nucleus accumbens). In both CON groups (RAC and FAL), a two-community partition subdivided the striatum into nucleus accumbens and anterior putamen in one community and caudate and posterior putamen into another. Striatal circuitry is typically split into three functional domains: limbic (reward and reinforcement) involving the ventral striatum, associative (cognitive and executive function) involving the central striatal areas, and motor (sensorimotor control) involving the dorsolateral regions (Iversen et al. 2010; Martinez et al. 2003; Mawlawi et al. 2001). These three subdivisions follow an anterior-posterior gradients with topographically organized parallel and integrative circuits with the cortex (Haber 2016; Haber et al. 2006). The community organization of RAC BP, which largely divides the anterior and posterior striatal covariance network nodes is generally consistent with these functional domains of the striatum.

Another potential factor that could contribute to the observed covariance is the distribution of DA D_2_/D_3_ receptors in the striatum. Both RAC and FAL are generally considered nonselective D_2_/D_3_ antagonists. While FAL has a higher D_3_ selectivity relative to RAC, it is not sufficient to consider it D_3_-selective (Doot et al. 2019). Autoradiography studies have shown that both D_2_ and D_3_ are present in the striatum, with greater D_2_ density throughout (Mach and Luedtke 2018). Additionally, there is evidence that D_2_/D_3_ density ratio differs in precommissural (anterior) and postcommissural (posterior) caudate and putamen (Sun et al. 2012). Taking this into account and in combination with the comparison of RAC and FAL covariance networks in Figure 5, it is plausible that regional receptor distribution contributes to the observed covariance.

After establishing that DAergic striatal covariance networks possessed organizational features, we investigated potential alterations in AUD, as substance use disorders are believed to disrupt DAergic system function (Koob and Volkow 2010). Our analyses showed that the covariance structure of the AUD group network is indeed altered, showing less organization and lower network degree, density, and strength (Figure 2), supporting previous literature that alcohol dependence disrupts dopaminergic function in the striatum (Spitta et al. 2023), while providing novel insight into the inter-regional relationships of DAergic function. We identified three edges in which covariance significantly differed between AUD and CON: ipsilateral left hemisphere nucleus accumbens shell and anterior caudate, and contralateral left accumbens core and right hemisphere accumbens shell and core (Figure 2A). There is prior evidence of asymmetry in alcohol induced dopamine release in the ventral striatum (Oberlin et al. 2015; Yoder et al. 2016) and therefore it may be that left-right synchrony of DA function is disrupted in chronic alcohol misuse.

Finally, we carried out an exploratory analysis where the sample was stratified into cigarette smoking and nonsmoking AUD and nonsmoking CON participants. With only three smoking CON participants, the sample was insufficient to generate a covariance network. As expected, nonsmoking CON group had the highest network density, while the density was greater in smoking AUD (dual substance use) relative to nonsmoking AUD (single substance use) (Figure 4 A-C). Interestingly, co-assignment, a property of community structure of the networks, showed less organization in the smoking AUD network relative to the nonsmoking group networks (Figure 4D-F). While these exploratory findings are intriguing, further work is needed in larger samples and in otherwise healthy cigarette smoking participants to help determine the degree to which network disruption in co-use of tobacco and alcohol is distinct from alcohol or cigarette use alone.

These findings demonstrate the utility of a network-based approach in studying alcohol and substance use disorders such as AUD. The covariance strategy has been long applied to fluorodeoxyglucose (Horwitz et al. 1984), amyloid, and tau (Franzmeier et al. 2020; Gonzalez-Escamilla et al. 2021) PET data. Recently, the emphasis on what has been referred to as ‘molecular connectivity’ has begun to grow (Hansen et al. 2022; Sala et al. 2023). It is important to consider that covariance network applications require a homogenous group/sample and enough participants to accurately estimate the networks (Veronese et al. 2019). Additionally, inference is made at the group level, that is, with the assumption that any observed differences in two networks are driven by the variables of interest, which in the present work are AUD and cigarette use. Lastly, these networks help us to understand the potential significance of inter-regional relationships. However, care must be taken when interpreting the results, as the biological meaning is not as apparent as in conventional PET neuroimaging analyses. Additional work is needed to fully appreciate the functional relevance of interregional relationships in the striatal DA system.

In summary, we showed that striatal dopaminergic function can be successfully investigated using inter-subject covariance. In addition, we demonstrated the presence of similar organization in two separate D_2_/D_3_ PET tracers. We then showed that this organization is disrupted in AUD, with preliminary evidence for greater disruption in comorbid alcohol and cigarette use. Network approaches to neurotransmitter PET data, recently termed ‘molecular connectivity’ offer a novel avenue for increasing our understanding of alcohol and substance use disorders by studying interactions of regional neurotransmitter patterns.

