## Supplemental Figures for "Intra-Striatal Dopaminergic Inter-Subject Covariance in Social Drinkers and Nontreatment-Seeking Alcohol Use Disorder Participants"

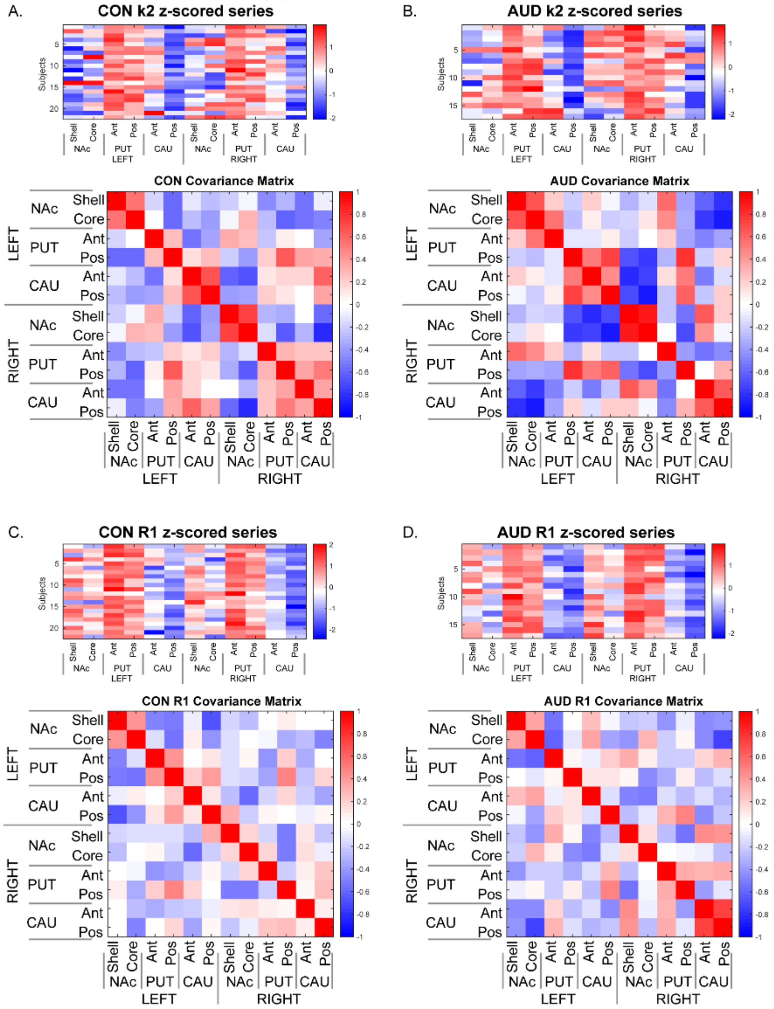


**Supplementary Figure 1.** *Z*-scored values of the tissue-to-plasma rate constant $k_2$ (A, B) and tracer delivery *R*1 (C, D) for social drinking controls (CON) and active alcohol use disorder (AUD) participants. Inter-subject covariance networks were computed as partial Pearson correlation, adjusting for age, biological sex, and smoking status. Ant – anterior, Pos – posterior, NAc – nucleus accumbens, PUT – putamen, CAU – caudate, CON – controls, AUD – alcohol use disorder.

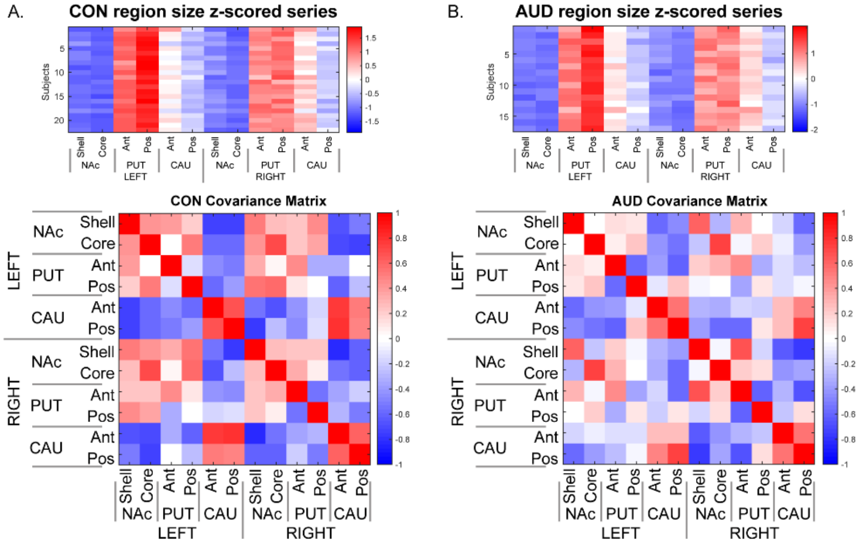


**Supplementary Figure 2.** Z-scored values and covariance networks of region size (in voxels) for social drinking controls (CON) and active alcohol use disorder (AUD) participants. Inter-subject covariance networks were computed as partial Pearson correlation, adjusting for age, biological sex, and smoking status. Ant – anterior, Pos – posterior, NAc – nucleus accumbens, PUT – putamen, CAU – caudate, CON – controls, AUD – alcohol use disorder.
